# Simultaneous dual recordings from vestibular hair cells and their calyx afferents demonstrate multiple modes of transmission at these specialized endings

**DOI:** 10.1101/2022.03.07.483321

**Authors:** Donatella Contini, Gay R. Holstein, Jonathan J. Art

**Affiliations:** Department of Anatomy & Cell Biology, University of Illinois College of Medicine, 808 S. Wood St, Chicago, IL 60612, USA; Neurology, Icahn School of Medicine at Mount Sinai, 1468 Madison Ave, New York, NY 10029, USA; Neuroscience, Icahn School of Medicine at Mount Sinai, 1468 Madison Ave, New York, NY 10029, USA

**Keywords:** synaptic transmission, hair cell, calyx, ion accumulation, quantal transmission, nonquantal transmission, ephaptic transmission, resistive coupling, vestibular

## Abstract

In the vestibular periphery, transmission via conventional synaptic boutons is supplemented by postsynaptic calyceal endings surrounding Type I hair cells. This review focusses on the multiple modes of communication between these receptors and their enveloping calyces as revealed by simultaneous dual-electrode recordings. Classic orthodromic transmission is accompanied by two forms of bidirectional communication enabled by the extensive cleft between the Type I hair cell and its calyx. The slowest cellular communication low-pass filters the transduction current with a time constant 10 — 100 milliseconds: potassium ions accumulate in the synaptic cleft, depolarizing both the hair cell and afferent to potentials greater than necessary for rapid vesicle fusion in the receptor and potentially triggering action potentials in the afferent. On the millisecond timescale, conventional glutamatergic quantal transmission occurs when hair cells are depolarized to potentials sufficient for calcium influx and vesicle fusion. Depolarization also permits a third form of transmission which occurs over tens of microseconds, resulting from the large voltage- and ion-sensitive cleft-facing conductances in both the hair cell and the calyx that are open at their resting potentials. Current flowing out of either the hair cell or the afferent divides into the fraction flowing across the cleft into its cellular partner, and the remainder flowing out of the cleft and into the surrounding fluid compartment. These findings suggest multiple biophysical bases for the extensive repertoire of response dynamics seen in the population of primary vestibular afferent fibers. The results further suggest that evolutionary pressures drive selection for the calyx afferent.

## Introduction

A persistent question in vestibular research is the functional significance of specialized calyx endings present on a subset of primary afferents in vestibular end organs. A majority of afferent and all efferent synapses involving hair cells (HCs) in the labyrinth and cochlea are mediated by conventional bouton endings. While the calyx represents a unique characteristic distinguishing vestibular from auditory sensory epithelia, the function of this morphologically specialized synapse remains obscure. Recent dual simultaneous recordings from HCs and their associated calyx afferents demonstrate that there are three modes of intercellular communication between them. In addition to unidirectional quantal transmission from HCs to afferents, there are two bidirectional forms of transmission. The slower of the two results from potassium flux into the cleft from either HC or afferent over tens of milliseconds, which can elevate [K^+^]_cleft_ from a resting ^~^8 to >28 mM. The corresponding E_(K)_ for channels facing the cleft are depolarized from −71 to −38 mV. The more rapid form of bidirectional transmission results from HC and afferent channels open near their respective resting potentials. These allow current flowing into the cleft from either HC or afferent to divide between a component flowing out of the cleft near its apex at the neck of the Type I HC and a component flowing across the cleft into the synaptic partner. As yet there is no compelling evidence that the calyx is uniquely associated with particular characteristics of afferent discharge regularity or dynamics.

## Anatomy, synaptic morphology, and innervation

Vestibular epithelia of amniotes contain two HC receptor subtypes. These were initially differentiated by HC cytology alone, and were subsequently defined by morphological differences in their associated afferent processes as well (1). Type I HCs are flask-shaped and are innervated by an afferent chalice surrounding the entire basolateral HC surface except the apical neck. The extensiveness and exclusivity of the cleft between the HC and its enveloping calyx, together with the tight junction seal protecting the endolymphatic space, creates a unique, diffusion-limited, femtoliter environment that is markedly different from perilymph. Type II HCs are cylindrical and participate in comparatively small synapses involving bouton-like endings of vestibular afferents that terminate on the receptor somata and the outer calyceal faces (2, 3).

Both vestibular HC subtypes utilize ribbon synapses for quantal transmission. These structures were originally recognized in retinal photoreceptors, where the ribbons were bedecked with synaptic vesicles, a few of which attached to the cell’s plasma membrane (4–8). Subsequent studies revealed variability in their overall structure (9, 10), resulting in the adoption of the more general term “synaptic body” to describe synaptic vesicles tethered to a central structure with a minority docked at an active zone.

Synaptic bodies have been reported in retinal photoreceptors and bipolar cells, pinealocytes, electroreceptors, and auditory and vestibular HCs (9). Both the morphology of the synaptic body, and the number, density, tethering and docking of the associated synaptic vesicles (10, 11) vary. In auditory HCs, this morphological diversity has been correlated with functional differences in peripheral auditory signal processing based on Ca^2+^-dependent exocytosis (12, 13). In both vestibular and auditory HCs, ribbon synapse morphology is thought to underlie the ability of HCs to respond with graded, high rates of sustained quantal transmitter release (14, 15). This dual function is achieved by the orchestrated availability of docked vesicles for immediate exocytosis, and tethered vesicles (the “readily releasable pool”) poised for imminent release (16, 17). As shown in Figure 1, ribbon synapses are present at Type I HC synapses with calyces, and Type II HC synapses with both boutons and calyx outer faces (2, 3, 18, 19). Efferent synapses with afferents and Type II HCs display conventional presynaptic membrane specializations and postsynaptic subsynaptic cisterns in the HCs. Despite this well-defined anatomy, however, the precise role of the calyx in shaping HC-afferent signaling is not known. In that context, it is noteworthy that Wersäll reportedly regretted his sequential naming of the HCs, deeming the Type I HC-calyx architecture more highly evolved (20).

**Figure 1.**
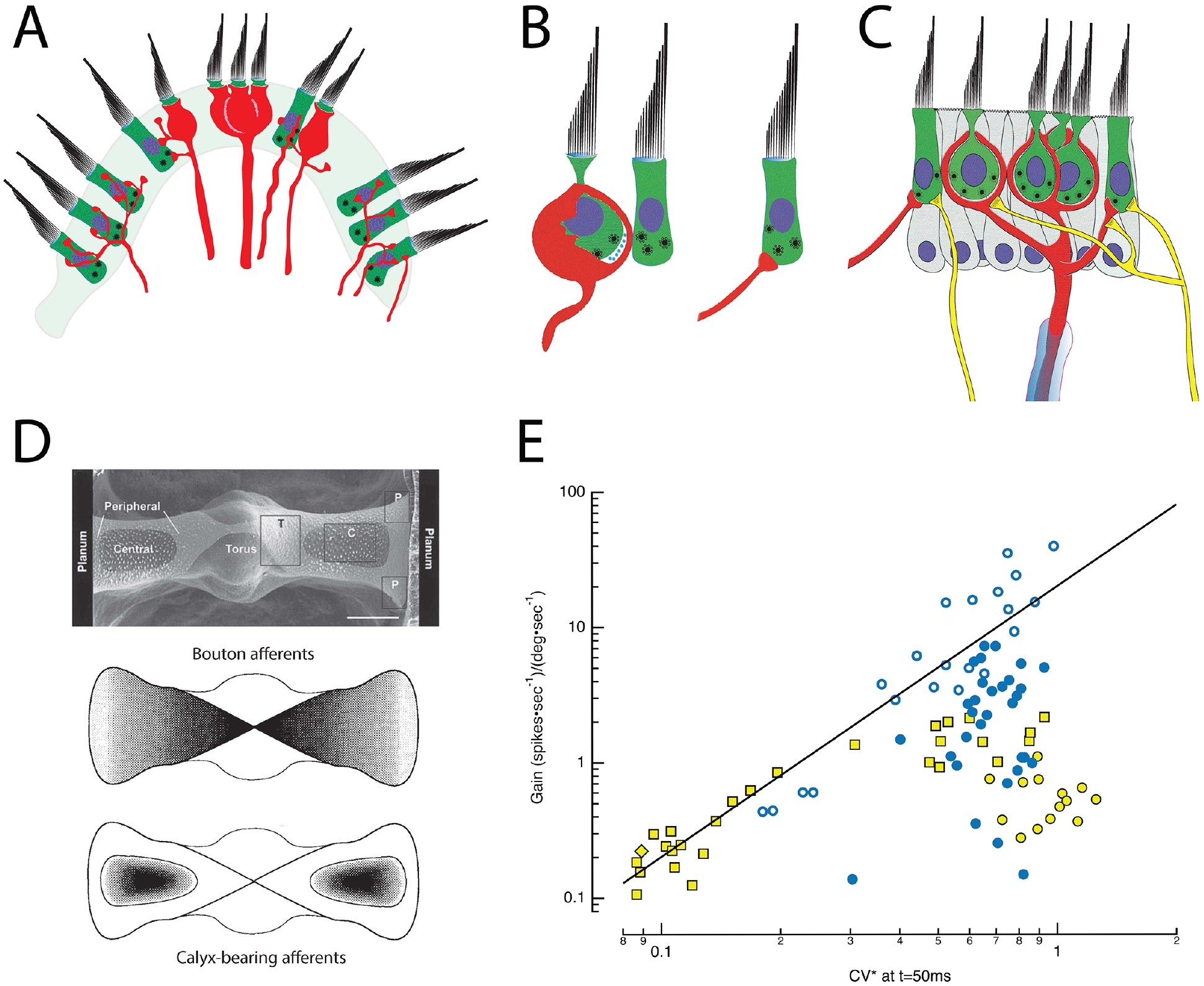
Morphophysiology of the semicircular canal (SCC) epithelium. A. Cross section through the saddle-shaped crista reveals distinct innervation patterns: boutons are found peripherally, with calyx and dimorphic afferents located centrally (adapted from (32)). B. Type I HCs are internal to an enveloping calyx that creates an extensive synaptic cleft in which ions accumulate. Type I HCs have synaptic bodies associated with quantal transmission onto the inner leaflet of the calyx. Type II HCs also have synaptic bodies indicative of quantal transmission, and synapse onto bouton endings (right) or onto the outer leaflet of an afferent calyx (middle and left). C. A single myelinated afferent (center, red) may branch into several calyces, each containing one or more Type I HCs. Additional input from Type II HCs may occur via synapses onto the outer face of the afferent (left), or through bouton endings of fine collateral branches (right). Efferent fibers (yellow) from the brainstem synapse on HCs and on the outer faces of calyces (60). D. Top: SEM plan view of the turtle posterior SCC (84). The epithelium is divided into two hemicristae with central and peripheral regions that extend bilaterally from the torus to the planum on either side. Middle: Map of the density of bouton afferents in each hemicrista. Bottom: Density map of the distribution of calyx-bearing afferents in the central regions of each hemicrista (36). E. Afferent signaling dynamics and discharge statistics in the posterior SCC of chinchilla (yellow symbols) (31) and turtle (blue symbols) (32) are correlated and map systematically across the epithelia. CV*, a measure of the irregularity of the interval between successive APs, is a continuous variable with the most regular fibers on the left and the most irregular on the right. Typically, when either animal was examined with low-frequency, 0.3 or 2 Hz, rotations, low gain afferents sensitive to velocity are more regular, and high gain afferents are sensitive to accelerations. For the chinchilla, the low gain bouton (yellow diamond) and dimorphic (yellow square) afferents demonstrate that regular discharge can be achieved with either bouton or calyx endings. Data seen on the right side, including high gain bouton (turtle: blue, open circle) and dimorphic (turtle: blue, filled circle; chinchilla: yellow square) fibers demonstrate that irregular discharge can be achieved by afferents containing either type of ending. The irregular low gain “calyx only” afferents in chinchilla (yellow filled circle) remain a puzzle but may simply be afferents that respond maximally to stimulations at higher frequencies (49) or with vibration (114). CV* for chinchilla data transformed from 15 ms inter-spike interval (ISI) CV* to that at 50 ms ISI using a third-order polynomial fit to the CV* vs ISI data at 50 ms in Fig. 1 of (31), equivalent to the CV* and ISI used in the turtle data (32).

## Morphophysiology

Decades of effort has focused on mapping morphological attributes of vestibular end organs and correlating these maps with dynamics and statistics of afferent firing (21–29); (Review: (30)). Based on intracellular recordings, dye injections and light microscopic reconstructions (31, 32), the crista ampullaris in multiple species has been subdivided into concentrically or linearly organized zones that evince differences in HC demographics and afferent innervation patterns (33, 34) (Fig. 1D). The posterior canal crista of turtles, for example, is organized into two symmetrical hemicristae separated by a central torus (35, 36). Each hemicrista has a central zone surrounded by a peripheral area that extends toward the torus and planum. All afferents in the central zone have calyces, manifest as either calyx-only fibers (32, 33) or dimorphic (37) fibers having both calyx and bouton endings. Afferents with bouton terminals are present in the peripheral zones throughout the crista, but more densely populate the zone with larger afferent fibers nearest the torus (32). Reflecting this, Type I HCs are more prevalent in the central zone while Type II HCs are present throughout the crista.

This differential localization of HCs and afferent endings across the crista underlies a coherent physiological and functional map of afferent activity (31, 33, 38–42). Specifically, the regularity of afferent fiber discharge, defined by the variability in time between sequential action potentials (APs) reportedly is a continuous variable correlating with the fiber’s response to velocity or angular acceleration (43–46). Regular afferents have evenly spaced spontaneous firing rates and respond to low frequency angular motion with tonic discharge. In effect, this mathematically integrates the acceleration stimulus, transmitting signals that are proportional to head velocity. In contrast, the spontaneous firing of irregular afferents shows high variability in the timing between APs. Angular acceleration elicits phasic, dynamic responses from these fibers, which more closely resemble the accelerative stimulus. Moreover, regularly discharging fibers as a group have lower gain (spikes*sec^−1^/degrees*sec^−1^) than irregular afferents. In turtles, low gain, regularly-firing bouton afferent endings sensitive to velocity are concentrated near the planum, while higher gain bouton afferents sensitive to acceleration terminate closer to the torus (47). In contrast, most calyx-bearing fibers, which end in the central zone, are irregular afferents. The sensitivity of these fibers to low-frequency rotation varies over two orders of magnitude and is not strictly correlated with the degree of irregularity of the spontaneous discharge rate. Nevertheless, an integrated morphophysiological map of the crista emerges in which low gain, regularly discharging, velocity-sensitive bouton afferents terminate nearest the planum, bouton afferents of higher gain and more irregular discharge terminate nearest the torus, and calyx-bearing, irregularly-firing units terminate in the central zone of each hemicrista (Fig. 1D). Functionally, the high gain fibers across species are particularly attuned to acceleration (47) or to jerks, the derivative of acceleration (48), and rapidly saturate as the amplitude of the stimulus increases. Medium gain afferents remain active over a wider motion spectrum, while low gain fibers are not prone to saturate, regardless of the magnitude of the head acceleration (Fig. 1E). One caveat to this characterization is that the afferent gains were typically assessed at single low frequencies of stimulation (31, 32), although the most irregular fibers are often maximally responsive to higher frequencies and even vibrational or sound stimuli (49–51).

Based on the initial correlation of afferent terminal distribution with regularity of spontaneous firing and response dynamics, significant effort was directed toward identifying causality. It remained to be determined whether the regularity of a fiber’s spontaneous firing derived from the geographical position of its terminals in the crista, or from the biophysical properties of HC inputs to trunk, branch, and twig segments of the parent process. Studies addressing this have been useful in highlighting the overall importance of regional localization in the crista and suggesting the type of HCs, three-dimensional terminal architecture and synapses contributing to individual afferent responses. However, efferent innervation (52–56), GABA-mediated modulation of glutamatergic transmission (57–59), ion accumulation in the synaptic cleft (60–63), and Type II HC input to the external face of the calyx (3, 64) have not been incorporated into these analyses. Since each of these has the potential to markedly alter peripheral vestibular activity, conclusions from these studies are necessarily tentative.

## Solitary cells

To supplement information gleaned by morphophysiology, currents were analyzed from HCs classified as Type I or II based on their amphoral or cylindrical profile (63, 65–83), from identified regions of the epithelia (84). This approach emulated a strategy used in lower vertebrates to demonstrate that the kinetics of voltage- and ion-sensitive basolateral channels imparted frequency selectivity to solitary auditory HCs (85–91). Typically, solitary vestibular HCs were bathed in artificial perilymph, an ionic environment appropriate for Type II cells, but of unknown propriety for the ionically dynamic milieu of a Type I HC and its calyx. In principle, such a bath would be apposite for isolating conductances based on their voltage ranges of activation, kinetics, ionic permeabilities, and pharmacological profiles. In artificial perilymph however, analysis of fundamental metrics such as I-V curves would be altered by driving forces reflecting equilibrium potentials for each ion when bathed in high sodium and low potassium. The vestibular potassium channels characterized in this way have been reviewed (92). A startling discovery was that Type I HCs possess a low-voltage outward-rectifying potassium current, IK_(LV)_, that activates at very hyperpolarized potentials. Its full activation near the HC resting potential was considered problematic, since enormous transduction currents would be required to depolarize a cell to potentials necessary for the calcium influx and vesicle fusion associated with quantal transmission. A solution to this paradox was proposed by Chen (93), who suggested that potassium accumulation in the synaptic cleft between the HC and calyx was analogous to the potassium accumulation in tissues surrounding the squid giant axon (94), and that potassium flowing from the HC would increase [K^+^]_cleft_ and depolarize E_K_ for HC and afferent conductances facing the cleft. As a result, the Type I HC would be depolarized at least to potentials necessary for calcium influx and quantal transmission, but the increased [K^+^]_cleft_ would also depolarize the calyx.

## Theoretical and single-electrode studies

Theoretical and experimental approaches were then used to examine the Chen conjecture. The theoretical analysis (95, 96) relied on equivalent circuits of the Type I HC, cleft, and calyx to analyze the problem primarily as ephaptic transmission (97–99) embellished by potassium accumulation. In general, this analysis concluded that the potential drops associated with current flow along the cleft from HC base to apex created only minor differences in the driving force for channels facing the cleft, and as a result were not likely to be a major source of coupling (96). The possibility that ion accumulation could have the effect suggested by Chen was also examined experimentally (100). Following blockade of quantal transmission, slow potentials that were phase-locked to the stimulus could be detected, a phenomenon consistent with ion accumulation. Two groups examined the potassium accumulation question by perforating the calyx and patch-recording from the enveloped Type I HCs *in situ* (101–103). These experiments demonstrated that depolarization of either the HC or the calyx caused a reversal of the HC potassium tail currents, suggesting an elevation of [K^+^]_cleft_ consistent with Chen’s conjecture. Such studies, together with one suggesting a transmitter role for protons (62), yielded results in which the interactions/coupling between cells occurred over tens of milliseconds. However, experiments using mechanical stimulation of the HC bundle demonstrated that when quantal transmission was blocked, the HC-afferent coupling could be extremely rapid, which would facilitate transmission of high frequency information (104, 105). Such contradictory viewpoints evoke the Indian parable of blind men describing an elephant (106), where each detects and interprets a small part of the more complex whole. This led to a quest to discover underlying biophysical mechanisms that could reconcile the differing results.

## Dual-electrode studies

The demonstration that [K^+^]_cleft_ can be increased by potassium fluxes from either synaptic partner is a reminder that interpretations of single-electrode voltage-clamp studies of either HC or afferent alone can be problematic. The privileged ionic compartment created by the apposition between a HC and calyx means that any single-electrode study that voltage clamped one cell necessarily allowed the potential of the other cell to vary freely. Moreover, studies that patched onto Type I HCs *in situ* (101–105) did so by perforating the calyx, and there is ample evidence that this approach compromises the cleft environment and results in two types of recordings: those similar to solitary cells bathed in perilymph, and those in which reversal of the potassium tail currents demonstrate an increase in [K^+^]_cleft_ (101, 102).

To address this, we developed an approach to simultaneously patch the HC and calyx without breaching the inner calyceal membrane (60, 61). In essence, this requires patching the HC either on its apex or on the exposed 2 – 3μm of its basolateral wall extending apically above the enveloping calyx. With this approach, the potentials of both cells can be interrogated or controlled in current clamp or voltage clamp, leaving the cleft ionic environment intact.

## Slow potassium accumulation depolarizes both HC and afferent to facilitate quantal transmission

Our initial studies (60) focused on ion accumulation and used the reversal potential of the HC potassium tail currents following depolarization to track E_K_ and estimate [K^+^]_cleft_. Blocking HC potassium flux abolished low-pass coupling of the transducer current occurring over tens of milliseconds, but spared quantal transmission. Our subsequent study (61) demonstrated that HC potassium flux depended upon the activation of both voltage-activated and calcium-activated potassium channels, and that the voltage-activated conductance was sensitive to [K^+^]_cleft_ and [Ca^2+^]_cleft_. Clamping [H^+^]_cleft_ resulted in stronger coupling between the two, suggesting that H^+^ was not a primary transmitter acting through acid-sensitive ion channels. Pharmacological experiments using an HCN channel blocker (60) demonstrated that, in addition to synaptic coupling acting through HC and afferent potassium channels, the elevated [K^+^]_cleft_ not only decreased outward K^+^ flux through calyceal voltage-sensitive potassium channels, but also shifted the activation curve of G_HCN_ (82), creating a depolarizing inward flux of Na^+^ and K^+^ through the HCN channels. These potassium-sensitive effects obtained regardless of whether the increased [K^+^]_cleft_ was due to HC or to afferent fluxes. Inward transduction currents would generate an outward flow of K^+^ through the basolateral HC, elevate [K^+^] and shift E_K_, causing depolarization of both HC and afferent (Fig 2A). Taken together, these results expand Chen’s conjecture that potassium accumulation in the cleft facilitates classical quantal transmission between a Type I HC and calyx, and extend it to direct stimulation of the afferent as well. For complex calyces with more than one enveloped HC, the findings indicate that depolarization of one HC can depolarize the afferent, which would then depolarize neighboring receptors enveloped by the shared afferent (Fig. 2A). In that light, slow potassium accumulation in the cleft, above that in perilymph, is the ongoing leaky integration of transduction currents or efferent excitation of the calyx, and thus can be interpreted as bidirectional inter-cellular communication that acts over tens of milliseconds.

**Figure 2.**
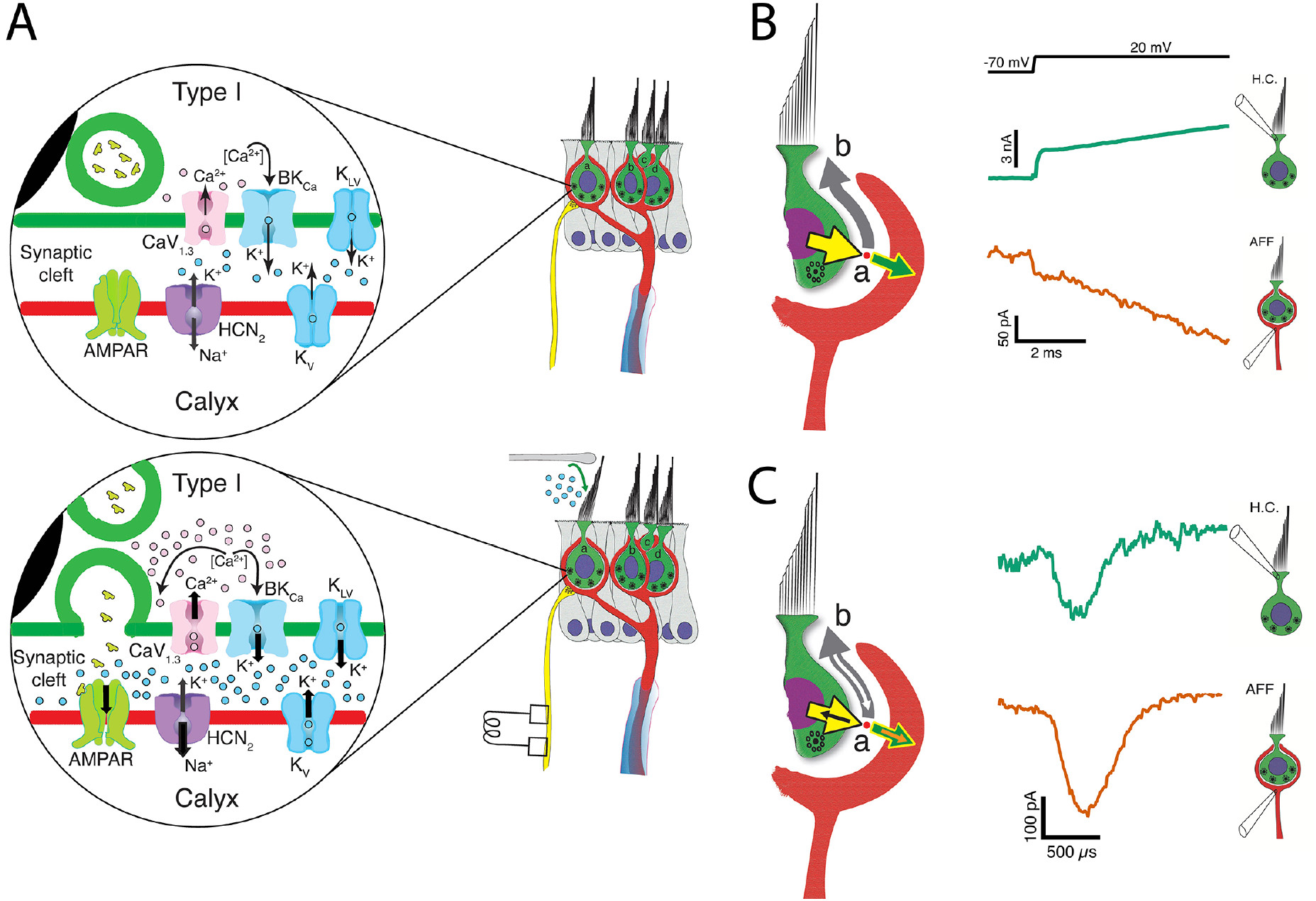
Three forms of synaptic transmission between a Type I HC and calyx afferent. A. In the absence of mechanical stimulation (upper panel), basolateral potassium currents activated at hyperpolarized potentials elevate the [K^+^]_cleft_ to concentrations near 8 mM, twice that found in perilymph. This results in a shift in E_K_ and depolarization of the HC to a resting potential between −60 and −50 mV, where Ca_1,3_ channels are activated. At rest there is occasional vesicle fusion and the production of a background rate of afferent discharge. Upon mechanical stimulation (lower panel), the inward transduction current flows into the HC and out the basolateral conductances into the cleft as a potassium current. The further increase in [K^+^]_cleft_ is a low-pass integral of the transduction current and further depolarizes E_K_ for conductances in the HC and afferent membranes facing the cleft. The HC depolarization gates Ca^2+^ influx, resulting in fusion of synaptic vesicles and quantal transmission of glutamate to AMPARs on the calyx inner leaflet. The slow accumulation of potassium thereby facilitates quantal transmission near the resting potential, and for maintained depolarizations towards 0 mV, elevates [K^+^]_cleft_ to greater than 28 mM within 25 ms. It is likely that the irregularities of vesicle fusion and large quanta that initiate an AP are driven by neighboring Ca-channel noise (118), and as a consequence the degree of regularity of afferent discharge would be determined by the degree to which the post-synaptic afferent conductances have intrinsic memory and oscillatory behavior similar to those found in HCs of the turtle auditory papilla (89, 119). B. Large conductances in the HC and calyx near the resting potential also create an open resistive pathway between the two that permits a fraction of the transduction current to be communicated across the cleft and into the calyx without synaptic delay. Outward current from the HC (yellow arrow) divides at point (a) into a large component (gray arrow) going along the cleft into the bath at point (b), and a smaller current flowing into the calyx (green arrow). The measured current for the HC (bluish-green current trace) and calyx (vermillion trace) show transmission within 10 μs. C. Resistive coupling can also be demonstrated during large HC depolarizations that depolarize the calyx to potentials sufficient to generate an AP that travels into the calyx. Currents due to the invasion of an AP will sum with the primary currents illustrated in B. With an AP, an additional inward current (orange arrow) flows into the calyx, with an amplitude equal to the sum of the corresponding currents flowing into the HC (black arrow) and down the cleft (white arrow). The coupling between the calyx retrograde AP current (vermillion) and the induced HC current (bluish-green) is 12 μs (modified from (61)).

## HC and afferent depolarization creates resistive coupling through increased open-probability of HC and afferent channels facing the cleft

One remarkable result of these experiments (61) was a demonstration that there is also a rapid bidirectional coupling between the cells that allows currents to activate within microseconds of depolarization. With depolarization near the HC and calyx resting potentials, there are a significant number of ion channels with high open probabilities. Rapid inward transduction current would immediately flow out as potassium through HC channels, with much of the current flowing into the extensive synaptic cleft and a minority flowing into the calyx (Fig. 2B). Such coupling can be analyzed using either Kirchhoff’s current or voltage laws, and the conclusion is the same: there would be relatively small voltage gains or drops of a few mV along the cleft depending on the current direction and the precise resistance (96, 107). The impact of this on specific channels facing the cleft is unknown and would greatly depend on their position with respect to the apical gap between HC and calyx. The dual recordings allow us to measure the currents flowing out of one synaptic partner and into the other, or out through the cleft (60, 61) (Fig. 2B, C). Such rapid current flow from HC into calyx, Fig. 2B, may underlie the rapid transmission of high frequency mechanical stimuli (104, 105, 108), and may be the substrate for detection of sound and vibration (26, 51, 109–115). As with slow potassium accumulation, this rapid ion flow through open channels is bidirectional, as evidenced by the retrograde transmission of calyx afferent APs into the HC (Fig. 2C) (60). Thus, as depolarization persists, [K^+^]_cleft_ increases, and both the number of open channels and the amount of HC-afferent electrical coupling increase concomitantly.

Dual recordings also provided the incidental observation that in the central region, the EPSCs from Type I HCs onto the calyx inner leaflet were small (60, 116, 117), whereas those from Type II HCs onto the outer leaflet (60) were as large as those reported in the cochlea (14), and would be sufficient to trigger APs in afferents near the spike threshold. This highlights the risk of morphophysiological studies defining calyx-only afferents based on light microscopy alone.

## Conclusions

During vertebrate evolution, selective pressures favored increased sensitivity and flawless encoding of three-dimensional vestibular information. This correlated with a higher prevalence of Type I HCs and calyx afferents (121, 122, 123, 134), which exhibit unidirectional quantal and two forms of bidirectional communication. Potassium accumulation in the synaptic cleft is a low-pass filter of HC transduction currents and afferent depolarizations by efferent fibers, and thus a leaky integrator of activity. At rest, [K^+^]_cleft_ is roughly 8 mM, or twice that found in perilymph, and is sufficient depolarize E_K_ and facilitate conventional quantal transmission. Maintained low-frequency excitation elevates [K^+^]_cleft_ further to >28 mM, a value sufficient to trigger afferent APs even in the absence of quantal transmission. From an evolutionary perspective, the pressure to extend transmission to higher frequencies was best served by direct resistive coupling created by large HC and afferent cleft-facing conductances present at and depolarized from rest. In evolving from bouton, through protocalyx to calyx, these conductances elevate [K^+^]_cleft_, depolarize E_K_ to facilitate quantal transmission, and provide a resistive path between synaptic partners that has minimal delay, is dynamic, and would increase as mechanical inputs become larger.

## Author contributions

All authors have made a substantial, direct, and intellectual contribution to the work and approved it for publication.

## Funding

This work was supported by NIH National Institute on Deafness and Other Communication Disorders grants R21 DC017292; R21 DC017577, R01 DC019953, and R01 DC008846

## Acknowledgements

We would like to thank Professor A. Brichta for graciously allowing us to modify, reuse, and repurpose his figures in Figure 1.

## Notes

### Competing Interest Statement

The authors have declared no competing interest.

### Summary of Updates

To clarify the integration of the transduction current by the potassium accumulation in the cleft & therefore the slow bidirectional communication is a valid means cellular, or synaptic transmission.

